# The surface of lipid droplets constitutes a barrier for endoplasmic reticulum residential integral membrane spanning proteins

**DOI:** 10.1101/2020.07.28.225391

**Authors:** Rasha Khaddaj, Muriel Mari, Stéphanie Cottier, Fulvio Reggiori, Roger Schneiter

## Abstract

Lipid droplets (LDs) are globular subcellular structures that mainly serve to store energy in form of neutral lipids, particularly triacylglycerols and steryl esters. LDs are closely associated with the membrane of the endoplasmic reticulum (ER), and are limited by a monolayer membrane of phospholipids harboring a specific set of proteins. Most of these proteins associate with LDs through either an amphipathic helix or a membrane-embedded hairpin motif. Here we address the question whether integral membrane spanning proteins could localize to the surface of LDs. To test this, we fused perilipin 3 (PLIN3), a mammalian LD-targeted protein, to ER resident proteins, such as Wbp1 (a N-glycosyl transferase complex subunit), Sec61 (a translocon subunit), and Pmt1 (a protein O-mannosyltransferase). The resulting fusion proteins localize to the periphery of LDs in both yeast and mammalian cells. This peripheral LD localization of the fusion proteins, however, is due to redistribution of the ER around LDs, as revealed by bimolecular fluorescence complementation between ER- and LD-localized partners in cells coexpressing the membrane-anchored perilipin. A LD-tethering function of PLIN3-containing membrane proteins was confirmed by fusing PLIN3 to the cytoplasmic domain of OM14, an outer mitochondrial membrane protein. Expression of OM14-PLIN3 resulted in close apposition of mitochondria and LDs. Taken together, these data indicate that the LD surface constitutes a barrier for ER-localized integral membrane spanning proteins.

## Introduction

Lipid droplets (LDs) are present in most cells, where they are discernible as globular structures. Adipocytes typically contain one large unilocular LD, whereas other cells, such as yeast may contain up to a dozen of distinct LDs that are relatively small in size. LDs are mostly composed of neutral lipids, particularly triacylglycerols (TAG) and steryl esters (STE). This hydrophobic core of neutral lipids is surrounded by a phospholipid monolayer that harbors a specific set of proteins, many of which function in neutral lipid metabolism, such as lipases or acyltransferases. The morphology of LDs is somewhat reminiscent of that of apolipoproteins and milk globules, but unlike these structures, LDs are not secreted. LDs serve to store metabolic energy, which is released upon beta-oxidation of fatty acids esterified to these neutral lipids. In addition, lipids stored in LDs can also serve as readily available building blocks for rapid membrane proliferation. Thus, LDs can buffer both an excess and a lack of fatty acids, and thereby provide a protective role in lipotoxicity. Hence, LD homeostasis is associated with multiple prevalent human diseases including obesity and atherosclerosis (Olzmann and Carvalho, 2018, Thiam et al., 2013, Thiam and Dugail, 2019, Walther et al., 2017).

The biogenesis of LDs is driven by the formation of neutral lipids by endoplasmic reticulum (ER)-residential acyltransferases. Accordingly, the current model postulates that LDs originate from the ER membrane, where the accumulation of neutral lipids leads to the formation of lens-like structures within the ER bilayer. Growth of these neutral lipid lenses drives the formation of nascent LDs, which eventually emerge from the ER as mature LDs (Murphy and Vance, 1999). In both yeast and mammalian cells, LDs appear to stay in close association with the ER membrane, allowing for the exchange of proteins and lipids between these two compartments (Jacquier et al., 2011, Kassan et al., 2013, Markgraf et al., 2014, Wilfling et al., 2013, Zehmer et al., 2009).

The transfer of proteins and lipids between the ER and LDs is likely to occur through membrane contact sites between the two compartments. These membrane contacts are maintained by ER resident integral membrane proteins, including seipins and fat storage-inducing transmembrane (FIT) proteins, which are both required for proper LD biogenesis and modulate LD size and abundance (Kadereit et al., 2008, Szymanski et al., 2007). Members of the seipin protein family form large oligomeric complexes at ER-LD contact sites, and harbor an ER luminal lipid binding domain (Sui et al., 2018, Yan et al., 2018). Seipins have been proposed to act as a diffusion barrier at ER-LD contacts to uncouple the lipid composition of the ER membrane from that of LDs, thereby contributing to the establishment of LD identity (Grippa et al., 2015, Salo et al., 2016). FIT proteins, on the other hand, which include human FIT1 and FIT2, and yeast Scs3 and Yft2, bind TAG and diacylglycerols (DAG) *in vitro* and possess acyl-CoA diphosphatase activity (Becuwe et al., 2020, Goh et al., 2015, Gross et al., 2011). Lack of FIT proteins leads to the formation of LDs that remain enclosed in the ER, suggesting that these proteins function to promote cytoplasmic emergence of LDs (Choudhary et al., 2015).

Proteins can target the surface of LDs either from the cytoplasmic space or through the ER membrane (Dhiman et al., 2020). Soluble proteins, such as the abundant LD scaffolding perilipins (PLINs) and their yeast homologue Pet10, are cytosolic in the absence of LDs, but are recruited onto the surface of LDs upon induction of their biogenesis (Brasaemle, 2007, Bulankina et al., 2009, Gao et al., 2017, Jacquier et al., 2013). PLINs contain 11-mer amphipathic repeat segments similar to those present in apolipoproteins and the Parkinson’s disease-associated alpha-synuclein. These amphipathic repeats are important for LD targeting (Copic et al., 2018, Gao et al., 2017, Rowe et al., 2016).

LD-localized membrane proteins, on the other hand, are first inserted into the ER membrane from where they then redistribute to LDs (Kory et al., 2016). In the absence of LDs, these proteins display uniform ER localization and biochemically behave as membrane proteins. They are characterized by a hairpin type of membrane topology (Jacquier et al., 2011, Kassan et al., 2013, Wilfling et al., 2013, Zehmer et al., 2009). However, how exactly such membrane-anchored proteins move from the ER bilayer membrane onto the phospholipid monolayer of LDs is not well understood.

Here, we address the question whether ER residential integral membrane spanning proteins can be targeted to the surface of LDs. To test this, we generated chimeras of two well-characterized ER membrane proteins, Wbp1, a subunit of the oligosaccharyl transferase glycoprotein complex, and Sec61, a subunit of the ER translocon, with perilipin, PLIN3, which contains the information to target a protein to LDs. The resulting fusion proteins display circular staining of the LD periphery in both yeast and mammalian cells. Localization of these reporter proteins to the LD periphery, however, is not due to bona-fide localization of the perilipin-fused membrane protein on the surface monolayer of LDs, but is due to repositioning of the ER membrane around LDs. This close juxtaposition between LDs and ER is induced by the expression of membrane-anchored perilipin, as indicated by bimolecular fluorescence complementation (BiFC) of split GFP fluorescence between interaction partners localized on LDs and the ER. Consistent with an LD-tethering function of PLINs, fusion of PLIN3 to a membrane protein of the outer mitochondrial membrane, OM14, induces close apposition of mitochondria with LDs. Taken together, these results indicate that integral membrane spanning proteins are restricted from moving from the ER bilayer onto the limiting membrane of LDs, supporting the notion that the LD surface imposes a barrier to ER residential membrane proteins.

## Results

### Targeting of integral membrane spanning proteins to the perimeter of LDs

To test whether ER residential integral membrane spanning proteins could be targeted to the surface of LDs, we chose two well-characterized ER resident proteins: Wbp1, a single spanning membrane protein and component of the oligosaccharyl transferase complex required for N-linked glycosylation of proteins in the ER lumen, and Sec61, a multispanning transmembrane protein and subunit of the ER translocon (Deshaies and Schekman, 1987, te Heesen et al., 1992). Wbp1 and Sec61 were fused to a cassette consisting of a fluorescent reporter protein, GFP, and perilipin 3 (PLIN3/TIP47), a mammalian LD-localized protein (Bulankina et al., 2009, Wolins et al., 2001) (Fig. 1A). PLIN3 has previously been shown to localize to LDs when expressed in yeast and to associate with LDs through its amphipathic helix (Jacquier et al., 2013, Rowe et al., 2016). The expression of the reporter constructs was placed under the control of a tetracycline regulatable promoter (Tet-Off) and expression was induced by depleting doxycycline from the medium during overnight growth of the cells (Garí et al., 1997). The subcellular localization of these fusion proteins was then analyzed by fluorescence microscopy. Both Wbp1-GFP-PLIN3 and Sec61-GFP-PLIN3 localized to punctate structures, that were also stained with Nile Red, a LD specific dye (Fig. 1B). Colocalization was extensive since more than 86% of all GFP positive punctate structures were also stained with Nile Red (Fig. 1C).

**Figure 1.**
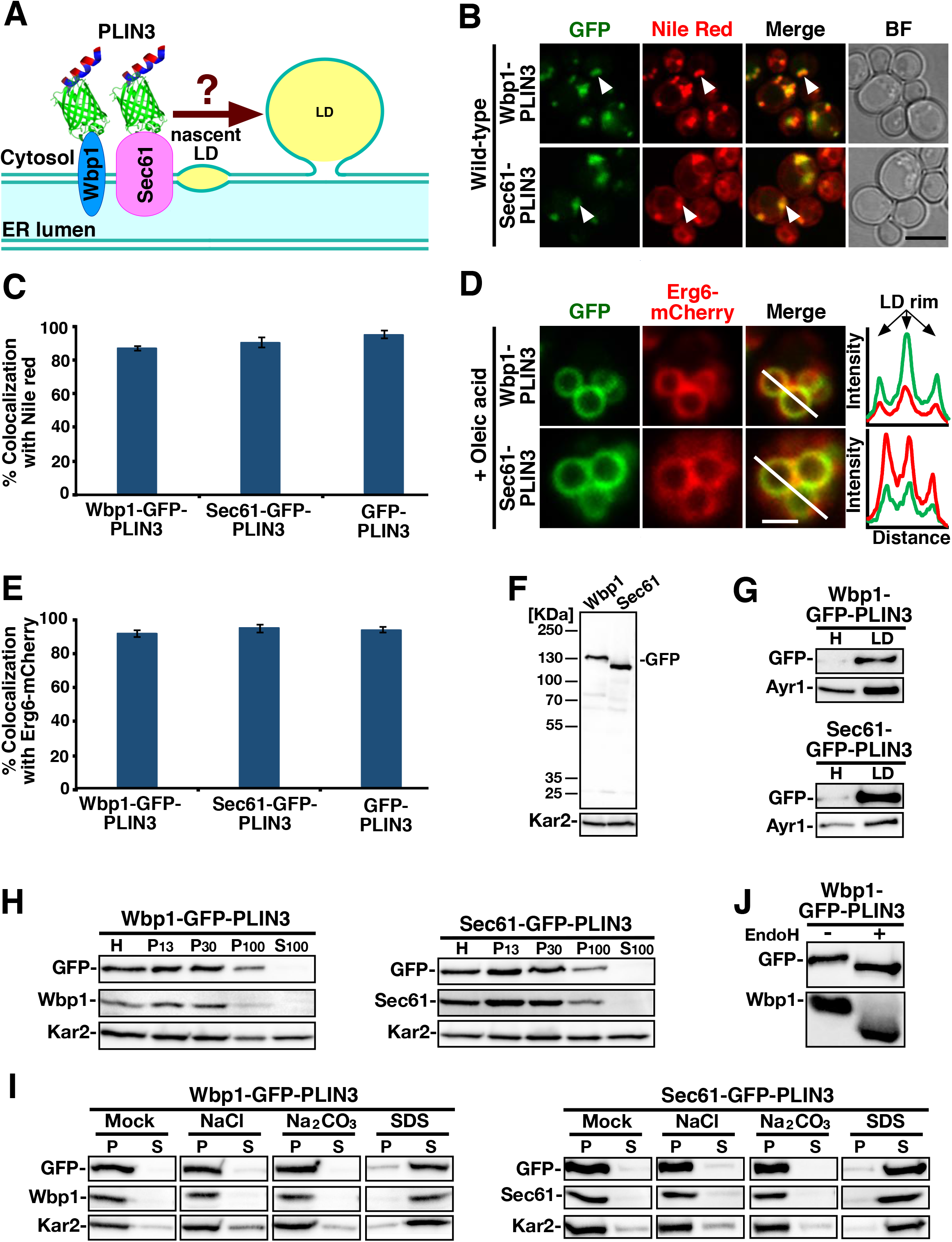
Membrane-anchored reporters localize to the periphery of LDs. A) Schematic structure of the membrane-anchored reporter proteins used in this study. The ER resident protein Wbp1 containing a single transmembrane domain, or the translocon subunit Sec61, containing multiple transmembrane domains, were fused to a reporter and targeting cassette consisting of GFP and perilipin 3 (PLIN3), a mammalian LD targeted protein. PLIN3 is indicated by the red/blue helix, GFP is represented by the green barrel, the transmembrane proteins, Wbp1 and Sec61, are shown by blue and pink ovals, respectively, and neutral lipids by yellow fills. B) Localization of Wbp1-GFP-PLIN3 and Sec61-GFP-PLIN3 to LDs in wild-type cells. Cells were grown in rich medium overnight at 30°C in the absence of doxycycline to induce expression of the reporters. Cells were stained with Nile Red and analyzed by confocal microscopy. White arrowheads indicate colocalization on LDs. BF, Bright field. Bar, 5 μm. C) Quantification of colocalization of Wbp1-GFP-PLIN3, Sec61-GFP-PLIN3, and GFP-PLIN3 with Nile Red stained LDs. N>100 LDs. D) Wbp1-GFP-PLIN3 and Sec61-GFP-PLIN3 colocalize with Erg6-mCherry at the periphery of LDs. Cells were cultivated in media containing oleic acid (0.12%) and the subcellular localization of -GFP and -mCherry tagged reporters was analyzed by fluorescence microscopy. A scan of fluorescence intensity along the white line that crosses two adjacent LDs is shown to the right. Bar, 2.5 μm. E) Quantification of colocalization of Wbp1-GFP-PLIN3, Sec61-GFP-PLIN3, and GFP-PLIN3 with Erg6-mCherry labelled LDs. N>100 LDs. F) Wbp1-GFP-PLIN3 and Sec61-GFP-PLIN3 are stable. Total cell extracts from wild-type cells expressing either Wbp1-GFP-PLIN3 or Sec61-GFP-PLIN3 were analyzed by Western blotting. The membrane was probed with antibodies against GFP and the ER luminal chaperone Kar2. G) Wbp1-GFP-PLIN3 and Sec61-GFP-PLIN3 are enriched on isolated LDs. LDs were isolated by flotation on step density gradients and an equal amount of protein (10 μg) from the homogenate (H) and the isolated LD fraction were probed by Western blotting using antibodies against GFP, and the LD-localized protein Ayr1. H) Wbp1-GFP-PLIN3 and Sec61-GFP-PLIN3 cofractionate with membranes. Cells were fractionated by differential centrifugation and individual fractions were probed by Western blotting using antibodies against GFP and either native Wbp1 or Sec61. H, homogenate; P13, 13,000 *g* pellet; P30, 30,000 *g* pellet; P100, 100,000 *g* pellet; S100, 100,000 *g* supernatant. I) Wbp1-GFP-PLIN3 and Sec61-GFP-PLIN3 become solubilized by detergent treatment only. Microsomal membranes were treated with salt (1 M NaCl), carbonate (0.1 M), or SDS (1%) for 30 min on ice, proteins were separated into membrane pellet and supernatant fraction, TCA precipitated, and probed with antibodies against GFP, Kar2, and Wbp1 or Sec61 to detect the native proteins. J) Wbp1-GFP-PLIN3 is N-glycosylated. Microsomal membranes were treated or not with endoglycosidase H (EndoH), TCA precipitated and analyzed by Western blotting using antibodies against GFP and Wbp1 to detect Wbp1-GFP-PLIN3 or the native version of Wbp1.

We then grew cells in the presence of oleic acid to increase the number and size of LDs. Both reporter constructs, Wbp1-GFP-PLIN3 and Sec61-GFP-PLIN3 localized to the periphery of LDs and perfectly colocalized with the LD marker protein Erg6-mCherry (Fig. 1D). This colocalization pattern was observed in more than 90% of LDs (Fig. 1E). Decoration of the LD periphery by these chimeras was not due to proteolytic cleavage of the GFP-PLIN3 reporter from the membrane anchor, as the fusion proteins were stable when analyzed by Western blotting and no major degradation intermediates could be detected (Fig. 1F). These data indicate that integral membrane proteins can be localized to the periphery of LDs.

To substantiate this result, we subsequently performed a membrane flotation assay. Wbp1-GFP-PLIN3 and Sec61-GFP-PLIN3 chimeras were enriched in the subcellular fraction containing LDs obtained by flotation, as was Ayr1, an LD marker protein (Athenstaedt and Daum, 2000) (Fig. 1G). Subcellular fractionation also indicated that these two constructs remained membrane-anchored despite their fusion to GFP and PLIN3. They were enriched in microsomal membrane pellet fractions (P13 and P30) as were endogenous Wbp1 and Sec61 (Fig. 1H). Thus, we conclude that the appendage of PLIN3 as an LD-targeting tag to Wbp1 and Sec61, does not affect their property to fractionate with microsomal membranes. Importantly, both fusion proteins stayed membrane-anchored upon treatment of the microsomal membranes with 1 M salt or carbonate, but they became solubilized upon treatment with 1% SDS, displaying the same membrane association properties as did endogenous Wbp1 and Sec61 (Fig. 1I). In addition, treatment of microsomal membranes with endoglycosidase H, resulted in a mobility shift of both the Wbp1-based chimera and native Wbp1, suggesting that the chimeric protein is glycosylated in its large ER luminal domain (Fig. 1J). Altogether, these results show that the generated fusion proteins conserved the biochemical properties of integral membrane proteins and their original topology, despite their localization at the periphery of LDs.

### Wbp1-GFP-PLIN3 and Sec61-GFP-PLIN3 induce local ER redistribution adjacent to LDs

To examine whether the localization of Wbp1-GFP-PLIN3 and Sec61-GFP-PLIN3 to LDs alters somehow the morphology of this organelle, we analyzed cells expressing Erg6-GFP, or one of the two chimeras at the ultrastructural level using immune-electron microscopy. As expected, Erg6-GFP specifically localized on the perimeter of LDs, and these organelles display membrane contact sites with the ER (Fig. 2A). In cells expressing Wbp1-GFP-PLIN3, labelling was found on the surface of LDs but also within a vesicular-tubular network (Fig. 2B, red square). In addition, these cells displayed long stretches of ER in close proximity of LDs, covering >70% of the LD perimeter. This was a peculiarity of this strain as in Erg6-GFP- or Sec61-PLIN3-GFP-expressing cells, only 15-23% of the LD perimeter was in clear contact with the ER (Fig. 2D). LDs in cells expressing Sec61-GFP-PLIN3 displayed gold labelling over their limiting membrane and frequent contacts with both the ER and the vacuoles (Fig. 2C). These ultrastructural observations confirm the confocal microscopy results, i.e., the Wbp1-GFP-PLIN3 and Sec61-GFP-PLIN3 chimeras are specifically recruited on the periphery of LDs and, particularly in cells expressing Wbp1-GFP-PLIN3, they also enhance the presence of ER-LD contact sites. Such extensive ER-LD contact sites have previously been observed in cells cultivated in oleic acid-containing medium, which enhanced LD production at the expense of phospholipid synthesis, particularly that of phosphatidylinositol (Chorlay et al., 2019).

**Figure 2.**
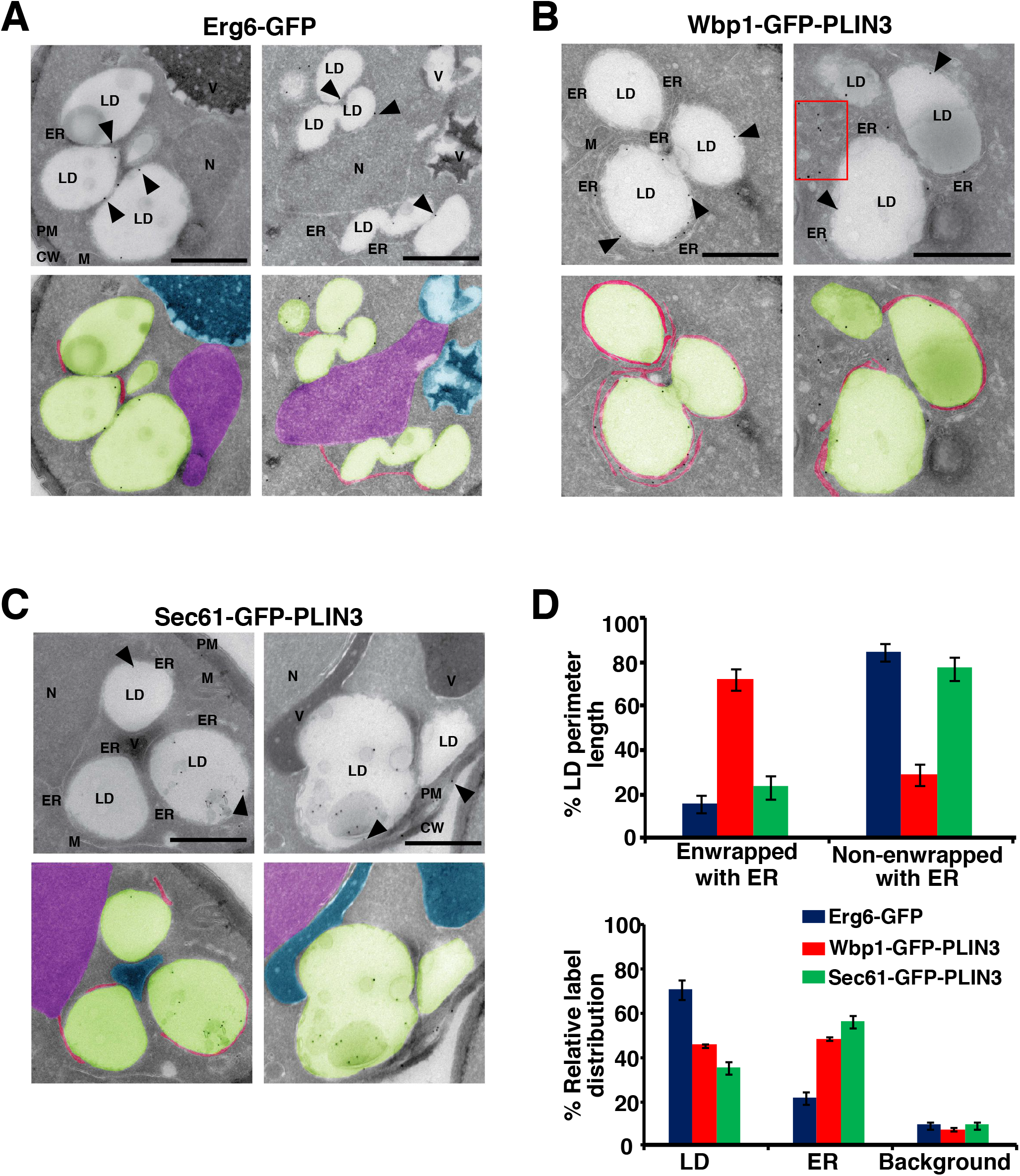
Immune-EM analysis of the subcellular distribution of the membrane-anchored LD reporters. A, B, C) Cells expressing Erg6-GFP (A), Wbp1-GFP-PLIN3 (B), or Sec61-GFP-PLIN3 (C) were grown in oleic acid supplemented medium (0.12%). Cells were collected, chemically fixed, processed for cryosectioning and immunogold labelled using an anti-GFP antibody and 10 nm protein A–gold particles indicated by black arrowheads. Organelles are indicated by colors: blue, vacuole; green, lipid droplet; pink, ER adjacent to LD; purple, nucleus. Red square, vesicular-tubular structures. CW, cell wall; ER, endoplasmic reticulum; LD, lipid droplet; M, mitochondria; N, nucleus; PM, plasma membrane; V, vacuole. Scale bar, 0.5 μm. D) Quantifications of the LD perimeter length enwrapped by the ER membrane and the relative distribution of the respective marker proteins between LDs and the ER.

### Integral membrane spanning proteins can be targeted to the perimeter of LDs in mammalian cells

Given that targeting of amphipathic helices to LDs is conserved among fungal, plant and animal cells, we tested whether mammalian ER membrane proteins could also be localized to LDs. Therefore, we created chimeric constructs composed of mammalian ER integral membrane spanning proteins fused to GFP-PLIN3. As ER membrane proteins, we chose OST48, a single spanning ER protein and a subunit of the mammalian oligosaccharyl transferase complex (Roboti and High, 2012), and SEC61A1, a multispanning ER protein and component of the mammalian translocon (Görlich et al., 1992). When transfected into HEK293 cells, OST48-GFP-PLIN3, SEC61A1-GFP-PLIN3, and GFP-PLIN3 all localized to punctate structures that were stained by Nile Red (Fig. 3A). These three chimeras also colocalized with mCherry-PLIN2 at the periphery of LDs (Fig. 3B). Colocalization of the chimeric fusion proteins with either Nile Red or mCherry-PLIN2 was typically >81%, irrespective of whether the cells were treated with oleic acid or not (Fig. 3C, D). Taken together, these data indicate that integral ER membrane spanning proteins can be targeted to the LD periphery in mammalian cells as well.

**Figure 3.**
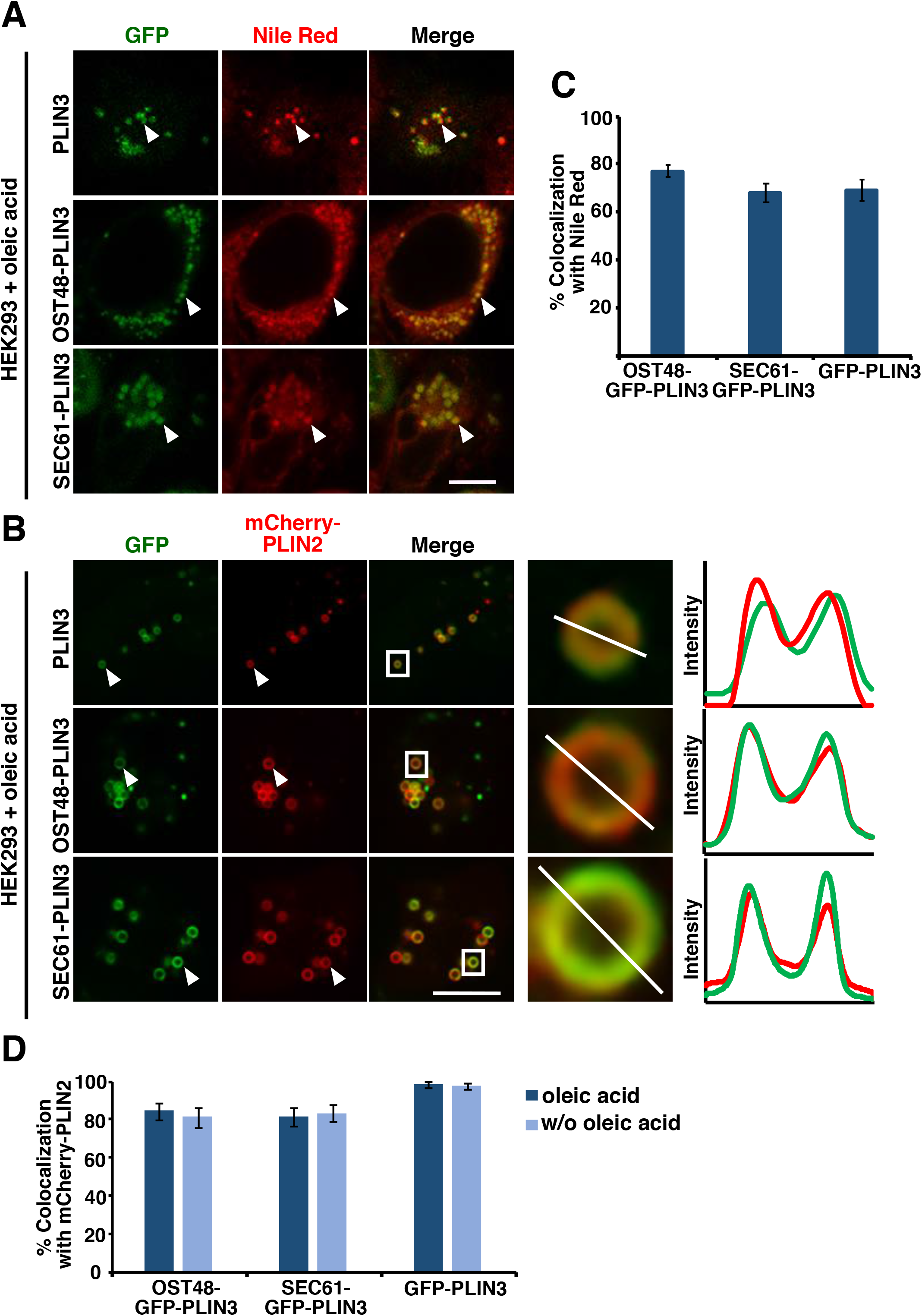
Integral membrane proteins can localize to the periphery of LDs in mammalian cells. A) Colocalization of soluble GFP-PLIN3, membrane-anchored OST48-GFP-PLIN3, and SEC61A1-GFP-PLIN3 with Nile Red stained LDs. Human embryonic kidney cells, HEK293, were transfected with the indicated GFP-tagged reporters, LDs were induced by cultivating cells in oleic acid containing medium for 4 h, and stained with Nile Red. LD labelling is indicated by arrowheads. Scale bar, 10 μm. B) Colocalization of soluble GFP-PLIN3, membrane-anchored OST48-GFP-PLIN3, and SEC61A1-GFP-PLIN3 with LDs labelled by mCherry-PLIN2. HEK293 cells were co-transfected with the indicated GFP-tagged reporters and mCherry-PLIN2, LDs were induced by cultivating cells in oleic acid containing medium for 4 h. Droplets marked by white arrowheads are boxed in the merge and enlarged in the panels to the right. Fluorescent intensity along the white line across the boxed LDs is shown in the graph. Scale bar, 10 μm. C, D) Quantification of colocalization of OST48-GFP-PLIN3, SEC61A1-GFP-PLIN3, and GFP-PLIN3 with LDs stained either by Nile Red (C) or labelled with mCherry-PLIN2 (D) in the presence or absence (w/o) of oleic acid. N>100 LDs.

### Appending PLIN3 to the ER-luminal side of an integral ER membrane protein fails to efficiently target the fusion protein to LDs

Given that appendage of PLIN3 to the cytosolically oriented end of integral ER membrane proteins such as Wbp1 or Sec61 conveys them to the periphery of LDs, we wondered whether the LD-localization of these reporters could be explained by redistribution of the ER around LDs rather than actual localization of the reporters on the limiting membrane of LDs, as schematically depicted in Fig. 4A. Such an ER redistribution-dependent localization of the reporters to the LD periphery, however, could occur only if the PLIN3 domain is exposed to the cytosol, as is the case with the two reporters studied so far. We thus tested whether fusion of PLIN3 to the luminal domain of a multispanning ER membrane protein, would still result in LD localization of the reporter. LD localization of a membrane reporter containing such an ER luminal PLIN3 could not be explained by ER repositioning. To test this, we chose to fuse GFP-PLIN3 to the C-terminus of Pmt1, a multispanning ER protein required for protein O-mannosylation, which has a topology that places its C-terminus in the ER lumen (Strahl-Bolsinger and Scheinost, 1999) (Fig. 4B).

**Figure 4.**
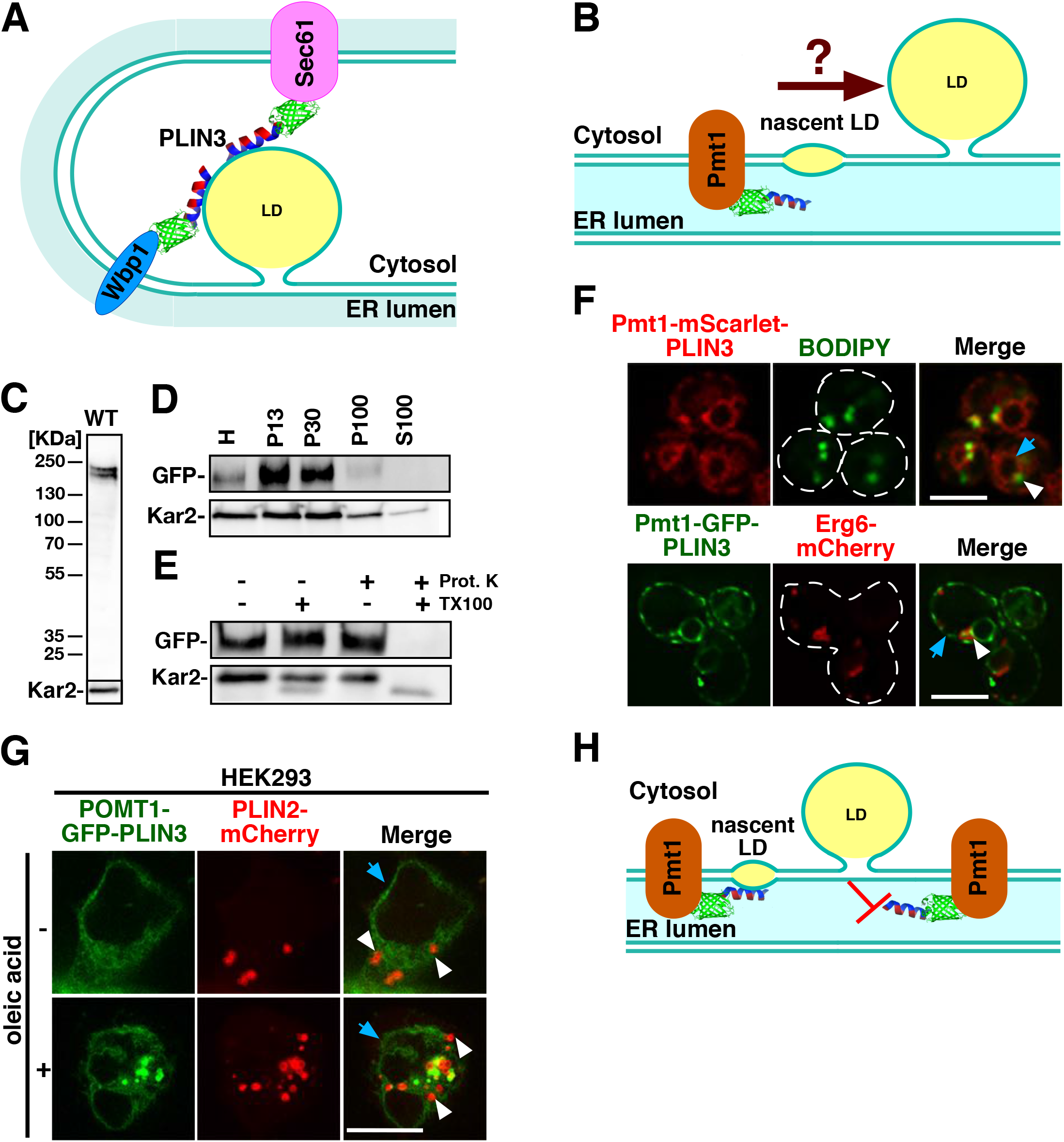
Localization of the LD targeting domain, PLIN3, in the ER lumen, fails to promote access to the LD periphery. A) Schematic representation illustrating the repositioning of the ER membrane around LDs, which could account for the observed LD localization of membrane-anchored reporters containing a cytosolic LD-targeting domain. B) Illustration of the topology of a Pmt1-based membrane-anchored reporter containing the LD targeting domain, GFP-PLIN3, within the ER luminal space. C) Full-length Pmt1-GFP-PLIN3 is stable. Western blot of total cell extracts from wild-type (WT) cells expressing Pmt1-GFP-PLIN3. The membrane was probed with antibodies against GFP and the ER luminal chaperone Kar2. D) Pmt1-GFP-PLIN3 fractionates with membranes. Cells expressing Pmt1-GFP-PLIN3 were fractionated by differential centrifugation and equal amount of proteins were probed by Western blotting using the ER luminal chaperone Kar2 as control. H, homogenate; P13, 13,000 *g* pellet; P30, 30,000 *g* pellet; P100, 100,000 *g* pellet; S100, 100,000 *g* supernatant. E) The GFP domain of Pmt1-GFP-PLIN3 is protected from proteinase K digestion. Microsomal membranes were treated with proteinase K (Prot. K) in the presence (+) or absence (−) of 0.1% Triton-X100 (TX100), proteins were TCA precipitated and probed with antibodies against GFP or Kar2. F) Pmt1-PLIN3 localizes to the ER membrane and not to LDs. Cells expressing Pmt1-mScarlet-PLIN3 were stained with BODIPY and analyzed by fluorescence microscopy. Cells coexpressing Pmt1-GFP-PLIN3 and Erg6-mCherry were grown in oleic acid containing media and analyzed by microscopy. ER localization of Pmt1-GFP-PLIN3 is indicated by the blue arrow, LD localization of Erg6-mCherry and the fluorescent dye BODIPY are highlighted by white arrowheads. Scale bar, 5 μm. G) POMT1-GFP-PLIN3 does not colocalize with PLIN2-mCherry labelled LDs in mammalian cells. HEK293 cells were cotransfected with POMT1-GFP-PLIN3 and mCherry-PLIN2, incubated in media supplemented with or without oleic acid for 4 h and imaged. ER staining by POMT1-GFP-PLIN3 is indicated by blue arrows. LDs are highlighted by white arrowheads. Scale bar, 10 μm. H) Illustration of the topological restrictions for LD localization of the ER luminal LD-targeting domain in Pmt1-GFP-PLIN3.

Pmt1-GFP-PLIN3 was expressed as full-length protein, fractionated with microsomal membranes and the GFP-PLIN3 domain was properly localized to the ER lumen as GFP was protected from protease digestion (Fig. 4C-E). When the subcellular distribution of Pmt1-mCherry-PLIN3 was analyzed by fluorescence microscopy, we found that this chimera localized to the ER membrane, and rarely colocalized with BODIPY stained LDs (Fig. 4F). ER localization of Pmt1-GFP-PLIN3 was also observed in cells coexpressing Erg6-mCherry grown in the presence of oleic acid (Fig. 4F). ER luminal exposure of an LD-targeting determinant is thus not able to target a protein to LDs. A membrane anchored perilipin thus behaves differently to a soluble GFP-PLIN3 reporter that localizes to LDs when targeted into the ER lumen (Mishra et al., 2016).

We next checked whether a membrane-anchored ER luminal PLIN3 would localize to LDs in mammalian cells. To this end, we appended GFP-PLIN3 to POMT1, a human O-mannosyltransferase orthologue of yeast Pmt1 (Jurado et al., 1999). When transfected into HEK293 cells, POMT1-GFP-PLIN3 colocalized to the ER and did not colocalize mCherry-PLIN2 on LDs (Fig. 4G). Thus, an ER luminal orientation of the LD-targeting domain does not promote the redistribution of an integral ER membrane spanning proteins to the periphery of LDs in both yeast and mammalian cells as illustrated in Fig. 4H. These results thus indicate that LD localization of membrane proteins containing a cytosolically oriented PLIN3 could be due to the repositioning of the ER membrane around LDs rather than *bona fide* localization of the reporter on the limiting membrane of the LD.

### >FITs affect access of the ER luminal PLIN3 to LDs

Next, we wondered whether the Wbp1-GFP-PLIN3, Sec61-GFP-PLIN3 and Pmt1-GFP-PLIN3 reporters can be targeted to mature pre-existing LDs. Therefore, we grew cells expressing Erg6-mCherry in oleic acid-containing medium to induce large LDs, but keeping the expression of the reporters repressed by the presence of doxycycline. Cells were then switched to medium lacking doxycycline to induce expression of the reporter and their localization was analyzed over time by fluorescence microscopy. Upon 2 h of induction, Wbp1-GFP-PLIN3 and Sec61-GFP-PLIN3, exhibited a crescent-like staining at the LD periphery (Fig. 5A). This crescent-shaped localization is likely to reflect the localization of the reporter at the ER-LD interface as has previously been observed for Gat1, a yeast glycerol-3-phosphate acyltransferase, which localized to crescent-like structures in the ER that are intimately associated with LDs (Marr et al., 2012). Similarly, the ER luminal Apolipoprotein B (ApoB) was localized to crescent-shaped areas adjacent to LDs in hepatoblastoma cells (Ohsaki et al., 2006). Upon overnight induction of the two membrane-anchored PLIN3 reporters, this crescent-like LD staining was superseded by a more uniform circular staining of the LD perimeter, as we previously observed in cells grown in the presence of oleic acid (Fig. 1D, 5A).

**Figure 5.**
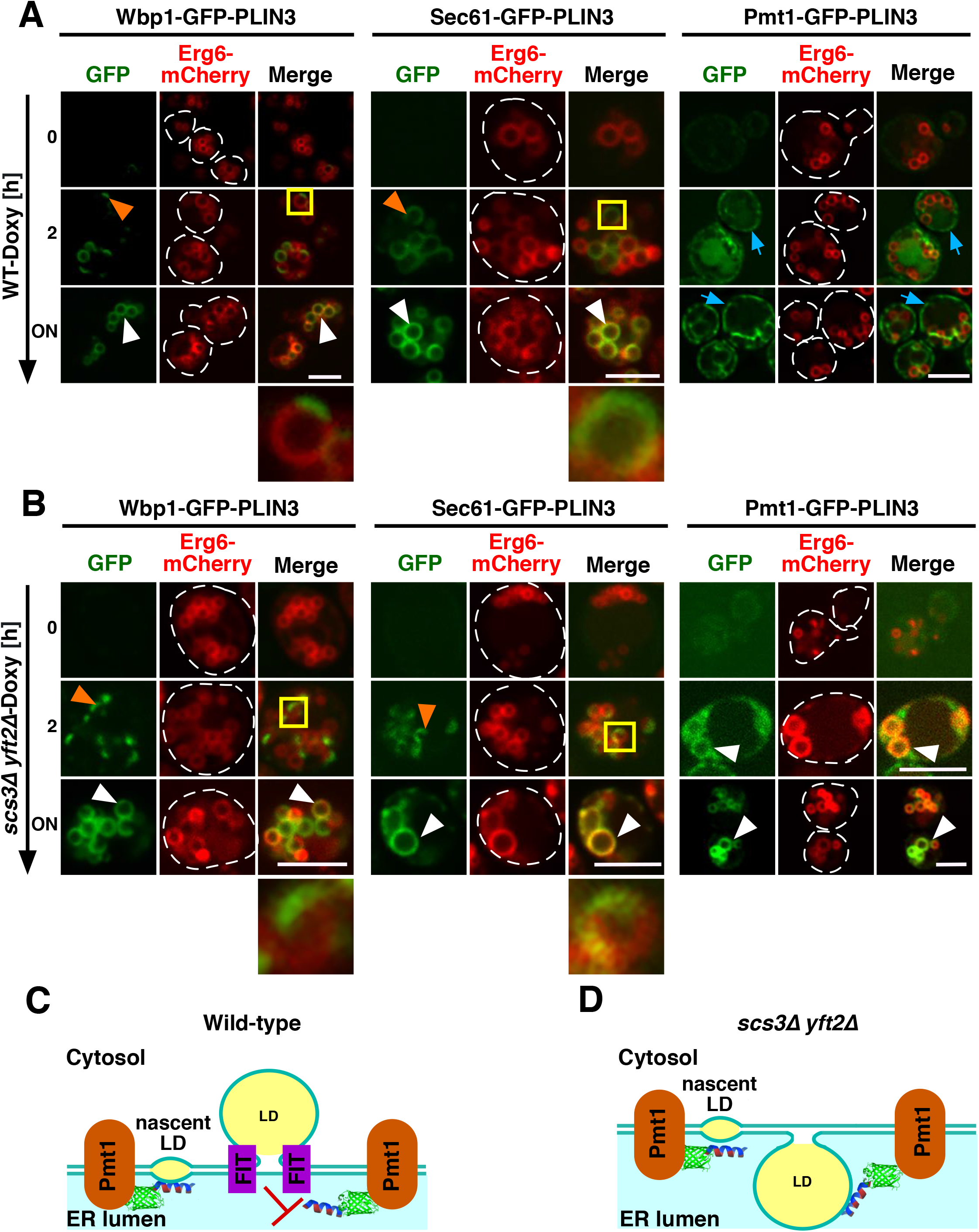
FIT proteins control access of an ER luminal targeting domain to LDs. A, B) Localization of membrane anchored reporters to pre-existing LDs. Wild-type and FIT double mutant (*scs3*Δ *yft2*Δ) cells expressing the indicated membrane-anchored LD reporters were cultivated in oleic acid supplemented medium, expression of the membrane proximal PLIN3 reporters was induced at time 0 by removing doxycycline (-Doxy) and their localization was analyzed by confocal microscopy. Orange arrowheads indicate crescent-like LD labelling of the reporters, white arrowheads depict the presence of the reporters around the LD periphery, and blue arrows mark the localization of Pmt1-GFP-PLIN3 in the peripheral ER. LDs boxed in yellow are magnified in the panels shown below. Note the circular staining at the LD periphery of Pmt1-GFP-PLIN3 in the *scs3*Δ *yft2*Δ double mutant, but not in wild-type cells. ON, overnight induction. Scale bar, 5 μm. C, D) Model to account for access of Pmt1-GFP-PLIN3 to mature LDs in FIT mutant cells. LDs emerging towards the cytosol as observed in wild-type cells are inaccessible to Pmt1-GFP-PLIN3, which contains the LD-targeting domain in the ER lumen. Emergence of LDs towards the ER luminal compartment in FIT mutant cells (*scs3*Δ *yft2*Δ) allows the access of Pmt1-GFP-PLIN3 to these luminally oriented mature LDs.

Induction of the reporter containing the ER luminally oriented PLIN3, Pmt1-GFP-PLIN3, in contrast, did not result in its targeting to pre-existing LDs (Fig. 5A). Instead, Pmt1-GFP-PLIN3 appeared to localize to the cell periphery, probably the cortical ER. These results indicate that the targeting of these membrane-anchored reporters to pre-existing LDs differs, depending on whether the LD targeting signal provided by PLIN3 has a cytosolic or an ER luminal orientation.

To analyze these differences in LD targeting in more detail, we examined the localization of these reporter constructs in FIT mutant cells. FIT proteins are required for the emergence of LDs towards the cytosol (Choudhary et al., 2015, Kadereit et al., 2008). The yeast genome encodes two FIT isoforms, Scs3 and Yft2. In the *scs3∆ yft2∆* double mutant, Wbp1-GFP-PLIN3 and Sec61-GFP-PLIN3 initially labelled LDs only partially, in a crescent-like manner, before dispersing over the entire LD perimeter, following the same kinetics as observed in a wild-type background (Fig. 5A, B). However, Pmt1-GFP-PLIN3 was localized to the circular perimeter of LDs in the FIT double mutant, even at the 2 h time point (Fig. 5B). These observations suggest that FIT proteins affect the access of membrane-anchored reporters to LDs. Thus, in the presence of functional FIT proteins, the membrane-anchored reporter containing an ER luminal LD-targeting domain, Pmt1-GFP-PLIN3, failed to properly localize to pre-existing LDs. However, in the absence of FIT protein function, Pmt1-GFP-PLIN3 could distribute to the perimeter of pre-existing LDs. The function of FIT proteins in controlling access of Pmt1-GFP-PLIN3 to mature LDs is redundant because the two single mutants, *yft2*Δ and *scs3*Δ, behaved as wild-type cells and prevented targeting of Pmt1-GFP-PLIN3 to pre-existing LDs (Fig. S1A, B). While FIT function affected targeting of Pmt1-GFP-PLIN3 to the perimeter of pre-existing LDs, it did not affect the stability of the protein, as revealed by a cycloheximide chase Western blot analysis (Fig. S1C). These observations support the hypothesis that FIT proteins affect the topology of LD budding, i.e., their emergence towards the cytosol or the ER lumen as depicted in the models shown in Fig. 5C, D (Choudhary et al., 2015). Emergence of LDs towards the ER lumen would thus allow the reporter with the ER luminal PLIN3 targeting domain, Pmt1-GFP-PLIN3, to gain access to pre-existing LDs.

### BiFC indicates that expression of the membrane-anchored PLINs induce repositioning of the ER membrane around LDs

To test whether the membrane anchored PLINs are in close proximity to established LD surface marker proteins, we employed a BiFC approach based on the split Venus readout (Magliery et al., 2005, Sung and Huh, 2007). Therefore, the N-terminal portion of Venus (-VN) was fused to the ER residents Wbp1 and Sec61, respectively, and the C-terminal part of Venus (-VC) was genomically fused to Erg6. Close proximity between the proteins fused to the two halfs of Venus, -VN and -VC, results in a BiFC signal generated by reconstitution of Venus (Fig. 6A). To test the functionality of this system, we first fused -VN to Erg6 and monitored reconstitution of fluorescence in cells coexpressing Erg6-VC. When grown in oleic acid containing medium, cells coexpressing Erg6-VN with Erg6-VC displayed BiFC on circular structures that completely colocalized with Erg6-mCherry, indicating that Erg6-VC is properly targeted to the surface of LDs and can thus be employed to monitor the proximity of VN-containing tester proteins (Fig. 6B). Next, we monitored BiFC in cells co-expressing Erg6-VC with the ER-residential Wbp1-VN or Sec61-VN. In both cases, Venus fluorescence was reconstituted in a crescent-shaped structure that localized to the periphery of LDs and partially overlapped with the circular fluorescence signal from Erg6-mCherry (Fig. 6C). These data thus indicate that the two ER proteins Wbp1-VN and Sec61-VN localized within a crescent-shaped domain of the ER membrane, at regions where the ER is in close proximity to LDs, to establish protein-protein interaction with Erg6-VC, as schematically drawn in Fig. 6D. In the absence of the VN-tag, both Wbp1-GFP and Sec61-GFP uniformly stain the ER membrane and did not accumulate at crescent-shaped ER-LD contacts (Fig. S2A).

**Figure 6.**
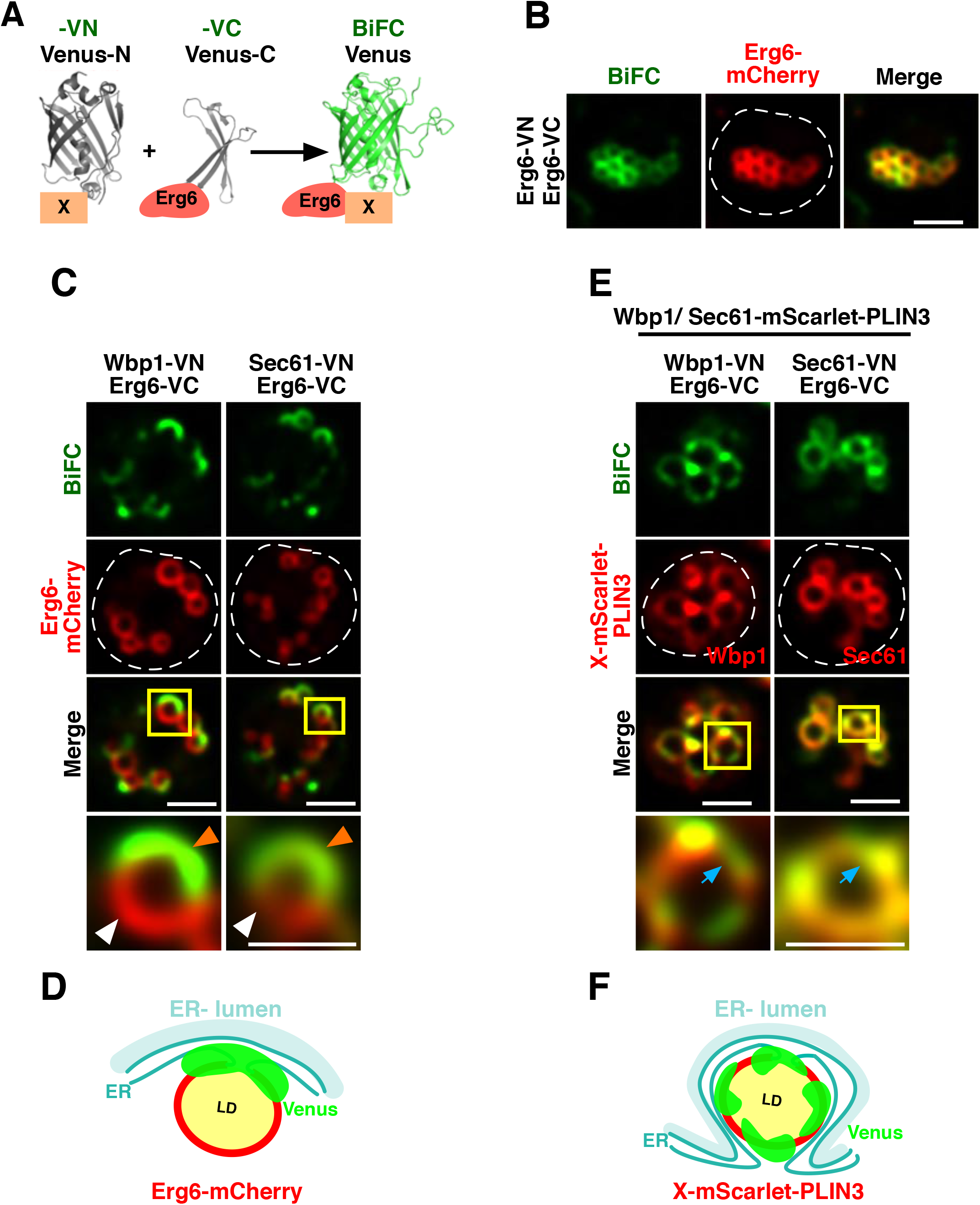
A split-Venus readout indicates that membrane anchored PLIN3 induces repositioning of the ER membrane around LDs. A) Schematic representation of bimolecular fluorescence complementation (BiFC) in which the N-terminal fragment of Venus (-VN) fused to a reporter protein X combines with the C-terminal fragment (-VC) from Venus fused to a second protein, in this case Erg6, to yield a fluorescent protein. B) Cells coexpressing Erg6-VC and Erg6-VN display BiFC on circular structures that colocalize with Erg6-mCherry at the periphery of LDs. Scale bar, 2.5 μm. C) Cells coexpressing Wbp1-VN and Erg6-VC or Sec61-VN and Erg6-VC display crescent-shaped labelling at the periphery of LDs. Cells expressing the indicated split-Venus fusions together with Erg6-mCherry were cultivated in oleic acid containing medium and analyzed by fluorescence microscopy. The regions enclosed by the yellow boxes are shown at higher magnification in the panels below. Note the crescent-shaped signal (orange arrowhead) from BiFC partially overlaps with the circular localization of Erg6-mCherry (white arrowhead). Scale bar, 2.5 μm. D) Illustration of the crescent-shaped BiFC signal representing contacts between the ER membrane, where Wbp1-VN and Sec61-VN are localized, and LDs, where Erg6-VC is localized. E) Expression of membrane-anchored PLIN3 extends the surface of interactions between Wbp1-VN or Sec61-VN and Erg6-VC to the entire LD perimeter. Cells expressing the indicated split-Venus fusions together with the membrane anchored PLIN3, Wbp1-mCherry-PLIN3 or Sec61-mCherry-PLIN3 were cultivated in oleic acid containing medium and analyzed by fluorescence microscopy. The regions enclosed by the yellow boxes are shown at higher magnification in the panels below. Note the circular signal from BiFC (blue arrow) overlaps with the circular localization of the mCherry-tagged membrane-anchored PLIN3. Scale bar, 2.5 μm. F) Illustration of the more circular-shaped BiFC signal representing extensive contacts between the ER membrane, where Wbp1-VN or Sec61-VN are localized, and LDs harboring Erg6-VC.

To test whether expression of the membrane-anchored PLIN3 would affect BiFC between Erg6-VC and either Wbp1-VN or Sec61-VN and thus potentially affect the ER-LD interface, we monitored BiFC in cells co-expressing red-fluorescent versions of the membrane-anchored PLIN3, Wbp1-mScarlet-PLIN3 or Sec61-mScarlet-PLIN3. While in the absence of the membrane-anchored PLIN3, the BiFC signal exhibits the above described crescent-shaped form, this was transformed to a more circular but nonhomogeneously distributed signal when cells co-expressed Wbp1-mScarlet-PLIN3 or Sec61-mScarlet-PLIN3 (Fig. 6E). This circular patched BiFC signal, however, overlapped with the localization of Wbp1-mScarlet-PLIN3 or Sec61-mScarlet-PLIN3, suggesting that expression of the membrane-anchored PLIN3 induced additional contacts between the ER and LDs as monitored by the BiFC signal in the green channel. Consistent with a close juxtaposition of the ER with LDs, Wbp1-GFP-PLIN3 and Sec61-GFP-PLIN3 colocalized with the ER luminal marker mCherry-HDEL in cells grown in the presence of oleic acid (Fig. S2B). These observations thus indicate that expression of the membrane-anchored PLIN3 promoted contacts between the ER and LDs along the LD circumference, suggesting that the ER membrane partially or completely enclosed the LD, as schematically illustrated in Fig. 6F. Taken together, these data indicate that the membrane anchored PLIN3, Wbp1-mScarlet-PLIN3 or Sec61-mScarlet-PLIN3, do not move from the ER bilayer onto the limiting membrane of LDs but that they induce the close apposition of the ER around LDs and thus induce redistribution of the ER around LD, resulting in a circular localization of the reporter around the LD perimeter.

### PLIN3 appended to an outer mitochondrial membrane protein induces close apposition of mitochondria with LDs

To test more directly whether membrane-anchored PLIN3 could induce close apposition of an organellar membrane with the surface of LDs, we fused PLIN3 to OM14, an outer mitochondrial membrane receptor for cytosolic ribosomes (Fig. 7A) (Lesnik et al., 2014). OM14-GFP colocalized with the mitochondrial marker Mito-RFP and LDs marked with soluble cytosolic PLIN3-GFP did not colocalize with mitochondria (Fig. 7B). However, when fused to PLIN3, OM14-GFP-PLIN3 colocalized with Nile Red stained LDs and when cells were grown in oleic acid supplemented medium, OM14-GFP-PLIN3 partially encircled Erg6-mCherry labelled LDs (Fig. 7C, D). In these cells, mitochondria started to surround LDs as indicated by the circular overlapping localization of Mito-RFP and OM14-GFP-PLIN3 (Fig. 7E). Close apposition of mitochondria with LDs in cells expressing OM14-GFP-PLIN3 was also observed by EM (Fig. 7F). From these results, we conclude that appendage of PLIN3 to a membrane protein can serve as an anchor to tether a membrane in close proximity to LDs, as illustrated in Fig. 7G. PLIN3 thus can serve as affinity tag to induce close apposition of an organellar membrane with LDs.

**Figure 7.**
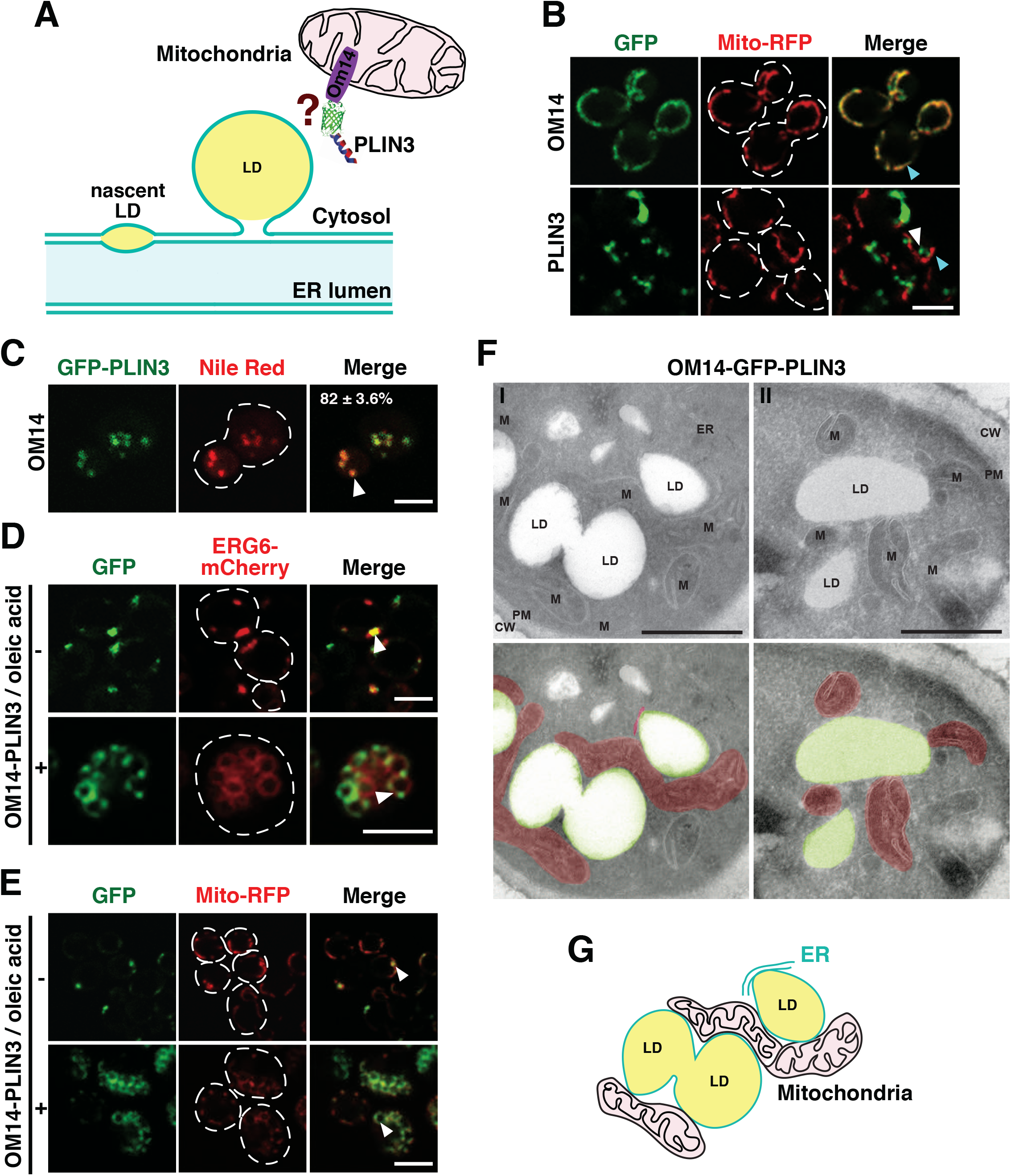
PLIN3 fused to an outer mitochondrial membrane protein induces close apposition between mitochondria and LDs. A) Schematic representation of the fusion of PLIN3 to OM14, an outer mitochondrial membrane protein. B) OM14-GFP colocalizes with the mitochondrial marker Mito-RFP but Mito-RFP does not colocalize with PLIN3. Cells coexpressing OM14-GFP with Mito-RFP or PLIN3-GFP with Mito-RFP were analyzed by fluorescence microscopy. Note the colocalization of OM14-GFP with Mito-RFP in the merge (turquoise arrowhead) whereas PLIN3-GFP (white arrowhead) does not colocalize with Mito-RFP. Scale bar, 5 μm. C) OM14-GFP-PLIN3 colocalizes with Nile Red stained LDs. Cells expressing OM14-GFP-PLIN3 were stained with Nile Red and their colocalization (highlighted by white arrowhead) was quantified, N>100 cells/LDs. Scale bar, 5 μm. D) OM14-GFP-PLIN3 colocalizes with Erg6-mCherry at the periphery of LDs. Cells coexpressing OM14-GFP-PLIN3 and Erg6-mCherry were cultivated in medium lacking (−) or containing (+) oleic acid and imaged by fluorescence microscopy. Colocalization is indicated by white arrowheads in the merge. Scale bar, 5 μm. E) LDs are in close proximity to mitochondria in cells expressing OM14-GFP-PLIN3. Cells coexpressing OM14-GFP-PLIN3 and Mito-RFP were cultivated in medium lacking (−) or containing (+) oleic acid and imaged. Colocalization is indicated by white arrowheads in the merge. Scale bar, 5 μm. F) Mitochondria are in close apposition with LDs in cells expressing OM14-GFP-PLIN3. Cells were cultivated in medium containing oleic acid, fixed and processed for EM. Organelles are indicated by colors: green, lipid droplet; red, mitochondria. CW, cell wall; ER, endoplasmic reticulum; LD, lipid droplet; M, mitochondria; N, nucleus; PM, plasma membrane. Scale bar, 0.5 μm. G) Schematic illustration of mitochondria positioning over the surface of LDs. LDs are indicated in yellow, mitochondria in pink, the ER membrane in green.

## Discussion

In this study, we address the question whether ER residential integral membrane spanning proteins can relocate over the limiting membrane of LDs and hence move from a membrane composed of a lipid bilayer into a lipid monolayer covering the hydrophobic core of LDs. Using protein fusions between ER membrane proteins and PLIN3, used as an LD targeting domain, we observe that these fusion proteins localize to the LD periphery and colocalize with a *bona fide* LD marker. However, this colocalization at the LD periphery is not due to actual relocalization of the ER residents to the limiting membrane of LDs, but caused by an apparent tight rearrangement of the ER membrane around the LD perimeter, as indicated by the BiFC results. These data thus indicate that regular transmembrane proteins cannot relocate over the lipid monolayer of LDs and hence that the limiting membrane of LDs effectively constitutes a barrier for ER residential membrane proteins. This is in contrast to membrane proteins containing a hairpin-type of membrane anchor, which can partition between the cytosolic leaflet of the ER membrane and the surface membrane of LDs and prefer to associate with a membrane monolayer rather than a bilayer *in vitro* (Caillon et al., 2020, Olarte et al., 2020). Whether the barrier for ER residential membrane proteins is established within the ER bilayer, i.e., at sites of contact between the ER and LDs, or whether it is the hemimembrane itself that prevents the repositioning of these membrane proteins from a bilayer membrane onto the monolayer remains an open question. The interface between the ER bilayer and the LD monolayer is constituted by both a seipin oligomeric ring and FIT proteins and is likely to contain additional factors such as LD associated factor (LDAF1/Promethin) and lipid modifying enzymes such as acyltransferases (Lro1) or the phosphatidate phosphatase (Pah1) and its regulators (Nem1/Spo7) (Choudhary et al., 2018, Choudhary et al., 2020, Chung et al., 2019, Salo et al., 2019, Yan et al., 2018). Whether and how hairpin-containing membrane proteins can pass through this protein interface or whether they are directly inserted into the limiting membrane of LDs remains to be established (Schrul and Kopito, 2016). Regular transmembrane proteins, however, are unlikely to pass through a seipin oligomeric ring, suggesting that the barrier for ER residential membrane protein is established within the ER itself.

The observation that expression of the membrane-anchored PLIN3 induces altered interactions between BiFC components localized in the ER and on the LD surface indicates that the expression of the membrane-anchored PLIN3 induces rearrangements of the ER membrane, probably redistributing the ER around LDs (Fig. 6E). Repositioning of the ER membrane is also observed by EM, particularly in cells expressing Wbp1-GFP-PLIN3, in which >70% of the LD perimeter is in close contact with the ER membrane (Fig. 2B). At the resolution of the fluorescence microscope, these ER-localized membrane-anchored PLIN3 then appear to completely colocalize with LD surface marker proteins, such as Erg6 (Fig. 1D). Membrane redistribution as the likely mechanism for the observed circular colocalization of the membrane-anchored PLIN with LD surface marker is further supported by the fact that PLIN3 appended to an outer mitochondrial membrane protein, OM14, induces close apposition between mitochondria and LDs (Fig. 7).

The observation that a membrane-anchored perilipin can induce tight apposition of a membrane such as the ER membrane or the outer mitochondrial membrane with LDs is unexpected and suggests that the interaction of PLIN3 with the LD surface is strong and persistent enough to induce membrane rearrangements. PLINs may thus serve as affinity tags to bring an organellar membrane in close proximity to LDs, which could be of interest for metabolic engineering, for example to increase the rate of fatty acid beta-oxidation by close apposition of mitochondria with LDs, a function that has been attributed to PLIN5 in oxidative tissue, especially heart (Wang et al., 2011). How exactly PLINs recognize and target the LD surface is not yet completely understood. PLINs contain repeats of amphipathic helices similar to those found on apolipoproteins, which also target a phospholipid monolayer that encloses a hydrophobic core of neutral lipids. These amphipathic helices have been proposed to recognize lipid packing defects on the lipid monolayer (Bulankina et al., 2009, Chorlay and Thiam, 2020, Copic et al., 2018, Prévost et al., 2018). The observation that induction of the PLIN3-containing membrane reporters in cells containing pre-existing large LDs initially results in their accumulation in crescent-shaped ER domains that are in close proximity to LDs, before they distribute over the entire LD surface, indicates that repositioning of the ER membrane around LDs is a slow process, which is consistent with a mechanism requiring membrane growth and repositioning (Fig. 5).

Membrane proteins containing an ER luminally localized LD-targeting domain, as is the case with Pmt1-GFP-PLIN3 in yeast and POMT1-GFP-PLIN3 in mammalian cells, fail to colocalize with LDs (Fig. 4F, G). This observation again supports the membrane redistribution-dependent mechanism for colocalization of PLIN3-containing membrane proteins with the LD periphery. However, this ER luminal LD-targeting domain can localize to pre-existing LDs in mutant cells lacking FIT proteins. FIT proteins and additional parameters such as surface tension, phospholipid synthesis and composition are important to impose a directionality to the LD budding process hence ensuring emergence of the LD towards the cytosol and not the ER lumen (Chorlay et al., 2019, Choudhary et al., 2015, Choudhary et al., 2018, Joshi et al., 2018, Wang et al., 2018). In the absence of FIT proteins or if surface tensions are altered, LDs can emerge towards the ER lumen, thereby topologically resembling the biogenesis of lipoproteins (Chorlay et al., 2019, Choudhary et al., 2015). The observation that membrane proteins containing a luminally oriented LD-targeting domain can localize to LDs in FIT mutant but not wild-type cells support the proposition that LDs bud towards the ER lumen rather than the cytosol in the absence of FIT function and provides for a readout to detect luminally oriented LDs. The previous observation that a soluble ER localized PLIN-GFP reporter can localize to LDs even in wild-type cells indicates that the soluble reporter can gain access to LDs under conditions where the membrane-anchored reporter is excluded from LDs and supports the existence of a membrane-localized barrier (Mishra et al., 2016).

Overall, the results of this study indicate that ER residential membrane proteins are excluded from the limiting membrane of LDs and hence that the LD surface could constitute a barrier for ER membrane proteins. This barrier may already be installed in the ER bilayer, for example in form of the seipin ring at the neck of LDs. How this seipin ring can be traversed by other membrane-anchored LD proteins, such as those containing a hairpin-type of topology will be interesting to understand in more detail.

## Acknowledgements

We thank all members of the lab for support, advice and helpful discussions, Vineet Choudhary and Aslihan Ekim Kocabey for comments on the manuscript, Mykhaylo Debelyy for analyzing protein topologies, Claire Jacob for advice on the construction of mammalian expression vectors and Mert Duman for preliminary localization experiments. F.R. is supported by ZonMW TOP (91217002), ALW Open Programme (ALWOP.310) and H2020 Marie Skłodowska-Curie Actions Cofund (713660) and Marie Skłodowska Curie ETN (765912) grants. M.M. is supported by an ALW Open Programme (ALWOP.355). R.S. is supported by the Swiss National Science Foundation (31003A_17303) and the Novartis Foundation for Medical-Biological Research (19B140).

## Materials and Methods

### Yeast strains, media and growth conditions

Yeast strains and their genotype are listed in supplementary Table S1. Yeast strains were cultured in YP-rich medium [1% bacto yeast extract, 2% bacto peptone (USBiological, Swampscott, MA)] or selective medium [0.67% yeast nitrogen base without amino acids (USBiological), 0.73 g/l amino acids)], containing 2% glucose. Fatty acid-supplemented medium contained 0.24% Tween 40 (Sigma-Aldrich, St Louis, MO) and 0.12% oleic acid (Carl Roth, Karlsruhe, Germany). To repress expression of the fusion constructs, cells were cultivated in the presence of doxycycline 1 μg/ml; Sigma-Aldrich, St Louis, MO). Expression was then induced by shifting cells to media lacking doxycycline, typically 14 h before imaging.

### Yeast expression constructs

Plasmid pCM189 containing the tetracycline-repressible *tetO*_*7*_ promoter (Garí et al., 1997) was used as a backbone to express the fusion proteins listed in the supplementary Table S2. The plasmid was linearized with BamHI and PstI, and a PCR fragment containing GFP-PLIN3 was inserted by homologous recombination in yeast (Hua et al., 1997), resulting in pCM189-GFP-PLIN3. pCM189-GFP-PLIN3 was then linearized by PacI and fragments encoding *WBP1*, *SEC61, PMT1,* or *OM14* PCR-amplified from yeast genomic DNA, were inserted by homologous recombination in yeast, resulting in Wbp1-GFP-PLIN3, Sec61-GFP-PLIN3, Pmt1-GFP-PLIN3, and OM14-GFP-PLIN3. To generate pCM189-mScarlet, the DNA encoding for mScarlet was codon optimized for expression in yeast and chemically synthesized (GenScript, Piscataway, NJ). mScarlet was then PCR amplified and cloned as a PstI/BamHI fragment into the pCM189 plasmid. Plasmids expressing Wbp1-mScarlet and Sec61-mScarlet were constructed by amplifying *WBP1* and *SEC61* followed by cloning into PacI-digested pCM189-mScarlet.

For construction of the split-Venus reporters, fragments encoding VN- or VC-terminal parts of Venus were amplified from plasmids pFA6a-VN-His3MX6 and pFA6a-VC-His3MX6, respectively (Sung and Huh, 2007). The plasmid pCM189 was linearized with BamHI and PstI, and a PCR fragment containing either Wbp1-VN, Sec61-VN, Erg6-VN, or Erg6-VC was inserted by homologous recombination, resulting in pCM189-Wbp1-VN, pCM189-Sec61-VN, pCM189-Erg6-VN, and pCM189-Erg6-VC respectively. All constructs were verified by sequencing (Microsynth AG, Buchs, Switzerland).

### Mammalian cell culture and expression constructs

Human embryonic kidney HEK293 cells were obtained from ATCC (Manassas, VA). Cells were cultured in DMEM (PAN-Biotech GmbH, Aidenbach, Germany) supplemented with 10% (v/v) fetal bovine serum (VWR International GmbH) and 0.2% penicillin-streptomycin (PAN-Biotech). Cells were seeded in 6-well plates and transfected the following day using lipofectamine 3000 (Thermo Fisher Scientific). 20 mM oleic acid complexed with 2.5 mM BSA were added to cells 16 h post transfection. Cells were incubated with oleic acid for 4 h prior to imaging. mCherry-PLIN2 expressed from the pLENTI6-mCherry-PLIN2 vector was used to label LDs (Eyre et al., 2014). SEC61A1-GFP-PLIN3 (Sino Biological, cDNA clone 19659-UT) and POMT1-GFP-PLIN3 (Sino Biological, cDNA clone 17179-UT) were cloned into pLENTI6 as NheI/FspI and NheI/ScaI fragments, respectively. OST48-GFP-PLIN3 (Sino Biological, cDNA clone 12463-UT) was cloned within the ScaI/NotI sites of pSiCoR-GFP-EF1α (Jacob et al., 2014). All constructs were verified by sequencing (Microsynth AG, Buchs, Switzerland).

### Protein fractionation and Western blot analysis

To analyse the subcellular distribution of the fusion proteins by fractionation, exponentially growing cells were harvested by centrifugation. Cells were then lysed in lysis buffer (20 mM HEPES pH 6.8, 150 mM potassium acetate, 250 mM Sorbitol, 1 mM DTT, 250 mM MgCl_2_, 2 mM PMSF and protease inhibitors (Roche Diagnostics, Mannheim, Germany)) by disruption with glass beads. The cell homogenate was cleared by centrifugation at 3,000 g for 5 min and then separated into pellet (P13) and supernatant (S13) fractions by centrifugation at 13,000 *g* for 30 min at 4°C. The S13 fraction was further separated into pellet (P30) and supernatant (S30) fractions by centrifugation at 30,000 *g* for 30 min at 4°C. The S30 was finally fractionated at 100,000 *g* for 1 h to yield a high-speed pellet (P100) and the cytosol (S100).

To determine the membrane association of the fusion proteins, 50 μg of proteins from the microsomal membrane pellet (P13) were incubated in lysis buffer containing either 1 M NaCl, 0.1 M Na_2_CO_3_, or 1% SDS for 30 min on ice. Samples were then centrifuged at 13,000 *g* for 30 min. Proteins in the pellet and the supernatant were precipitated with trichloroacetic acid (TCA) (10%), resuspended in SDS-PAGE loading buffer and analysed by Western blot.

For proteinase K protection experiments, 50 μg of proteins from the microsomal membrane pellet (P13) were incubated with 30 μg/ml of proteinase K (Roche Diagnostics, Mannheim, Germany) for 30 min on ice. Proteins were precipitated with TCA, and subjected to Western blot analysis.

For deglycosylation, 30 μg of microsomal proteins were treated with EndoH-Hf (New England Biolabs, Ipswich, MA) for 4 h at 37°C. The reaction was stopped by the addition of 1 mM PMSF and proteins were precipitated with 10% TCA. The pellet was dissolved in SDS-PAGE loading buffer and subjected to Western blot analysis.

LDs were isolated by two consecutive flotations as previously described (Leber et al., 1994). Briefly, spheroplasts were resuspended in lysis buffer (12% Ficoll PM 400, 10 mM MES-Tris, pH 6.9, 0.2 mM EDTA) and the lysate was cleared by centrifugation at 5,000 *g* for 5 min. This homogenate (H) was placed at the bottom of an ultracentrifuge tube, overlaid with lysis buffer and floated by centrifugation at 100,000 *g* for 1 h. The floating fraction was collected, diluted in lysis buffer and placed at the bottom of a second ultracentrifuge tube, which was overlaid with 8% Ficoll in 10 mM MES–Tris, pH 6.9, 0.2 mM EDTA and centrifuged at 100,000 *g* for 1 h. The second floating fraction corresponds to the crude LDs fraction. Proteins were delipidated with diethyl ether, TCA precipitated and analyzed by Western blotting. Protein concentration was determined by a Lowry assay, using the Folin’s reagent and BSA as standard.

GFP-fusion proteins were detected using a monoclonal antibody (Roche Diagnostics, diluted at 1:2000, #11814460001). Primary antibodies against Wbp1 (M. Aebi, ETH Zurich, Switzerland, diluted at 1:2000), Kar2 (R. Schekman, University of California, Berkeley, CA, diluted at 1:5000), Ayr1 (G. Daum, TU-Graz, Austria, diluted at 1:5000), or Sec61 (R. Schekman, diluted at 1:2000) were detected by using horseradish peroxidase (HRP)-conjugated secondary antibodies (Santa Cruz Biotechnology, Dallas, TX, diluted at 1:10,000, #sc-2030 and #sc-2302). Western blots and fractionation experiments were repeated at least two times with essentially similar results.

### Fluorescence microscopy

Yeast cells were grown to an early logarithmic phase (~1 OD_600nm_), pelleted by centrifugation and resuspended in a small volume of media. 3 μl of the cell suspension were mounted on a glass slide and covered with an agarose patch. Confocal images were recorded using a Leica TCS SP5 confocal microscope equipped with a 63x/1.20 HCX PL APO objective, and a LAS AF software. Cells were stained with Nile Red (10 μg/ml, Sigma-Aldrich, St Louis, MO) for 5 min at room temperature and washed twice with PBS. Localization of mCherry- and GFP-tagged fusion proteins was performed by fluorescence microscopy of live yeast cells using a Visitron spinning disk CSU-W1 (Visitron Systems, Puchheim Germany) or a DeltaVision deconvolution microscope (Applied Precision, Issaquah, WA). The Visitron spinning disk CSU-W1 consisted of a Nikon Ti-E inverted microscope, equipped with a CSU-W1 spinning disk head with a 50-μm pinhole disk (Yokogawa, Tokyo, Japan), an Evolve 512 (Photometrics) EM-CCD camera, and a PLAN APO 100x NA 1.3 oil objective (Nikon). The Delta Vision Elite (GE Healthcare, Pittsburgh, PA) imaging system consisted of an Olympus 1X71 inverted microscope equipped with a CCD camera (CoolSNAP HQ^2^, Photometrics, Tuscon, AZ). Images were acquired using a U PLAN S-APO 100x NA 1.3 oil immersion objective (Olympus). Eleven sections separated by 0.2 μm were deconvolved using the iterative constrained deconvolution program in softWoRx (Applied Precision, GE Healthcare, Pittsburgh, PA). Cells expressing the split-Venus constructs were imaged using the Visitron spinning disk microscope and images were deconvolved using Huygens Remote Manager, version 3.7.0 (http://svi.nl). Images shown are Z projection of 13-15 stacks collected at 0.25 μm step intervals.

Mammalian HEK293 cells were cultured in 6-well glass bottom plates (IBL, Gerasdorf, Austria). Cells were imaged at 16 h post-transfection with or without oleic acid-treatment using the Visitron spinning disk microscope. Images shown are Z projections of 20-30 stacks collected at a 0.5 μm step interval. Quantification of the GFP and mCherry signals was performed using the Fiji software (Schindelin et al., 2012). For Nile Red staining, cells were incubated with Nile Red (10 μg/ml; Sigma-Aldrich, St Louis, MO) for 5 min at 37°C, washed twice with PBS before fresh medium was added and images were collected using a Leica TCS SP5 (Leica, Wetzlar, Germany). A single confocal section is shown in the figures.

Colocalization was evaluated manually by scoring color overlap in 100 lipid droplets (% colabelling per cell). Images were treated using ImageJ software and then resized in Photoshop (Adobe, Mountain View). Microscopic experiments were performed three times with essentially similar results.

### Electron microscopy

For immune-electron microscopy, cells were grown in fatty acid supplemented medium at 30°C. Cell aliquots were collected by centrifugation, chemically fixed, embedded in 12% gelatin and cryosectioned as described previously (Griffith et al., 2008). Sections were then immunogold-labeled using a rabbit anti-GFP antiserum (Abcam, Cambridge, UK) and 10 nm protein A–gold, before being viewed in a CM100bio TEM (FEI, Eindhoven, Netherlands).

For the statistical evaluations, the number of LDs per cell section was determined by counting 100 randomly selected cell profiles. The relative distribution of the gold particles was calculated by classifying 550 of them, on the basis of their localization to the ER, LDs and mitochondria. A gold particle was assigned to a compartment when it was situated within 15 nm of the limiting membrane. The linear labeling density and the average organelle surface were established using the point hit method as described previously (Kondylis and Rabouille, 2003, Rabouille, 1999). The statistical significance of the data (*P*<0.05) was confirmed by *t*-test.

Quantification of ER-LD contacts was performed on three independent grids with a total of 30 random cell profiles for each genotype, using the intersection method (Rabouille, 1999). The relative distribution of the marker proteins was quantified by 4 independent evaluations of random cell profiles over three independent grids counting 682 gold particles for Erg6-GFP, 600 for Wbp1-GFP-PLIN3 and 659 particles for Sec61-GFP-PLIN3. Gold particles were assigned to a specific compartment, i.e., the ER or LDs, if they were positioned not more than 15 nm away from its limiting membrane (Mari et al., 2008).

## Supplementary Data

### Supplementary Figure Legends

**Figure S1.**
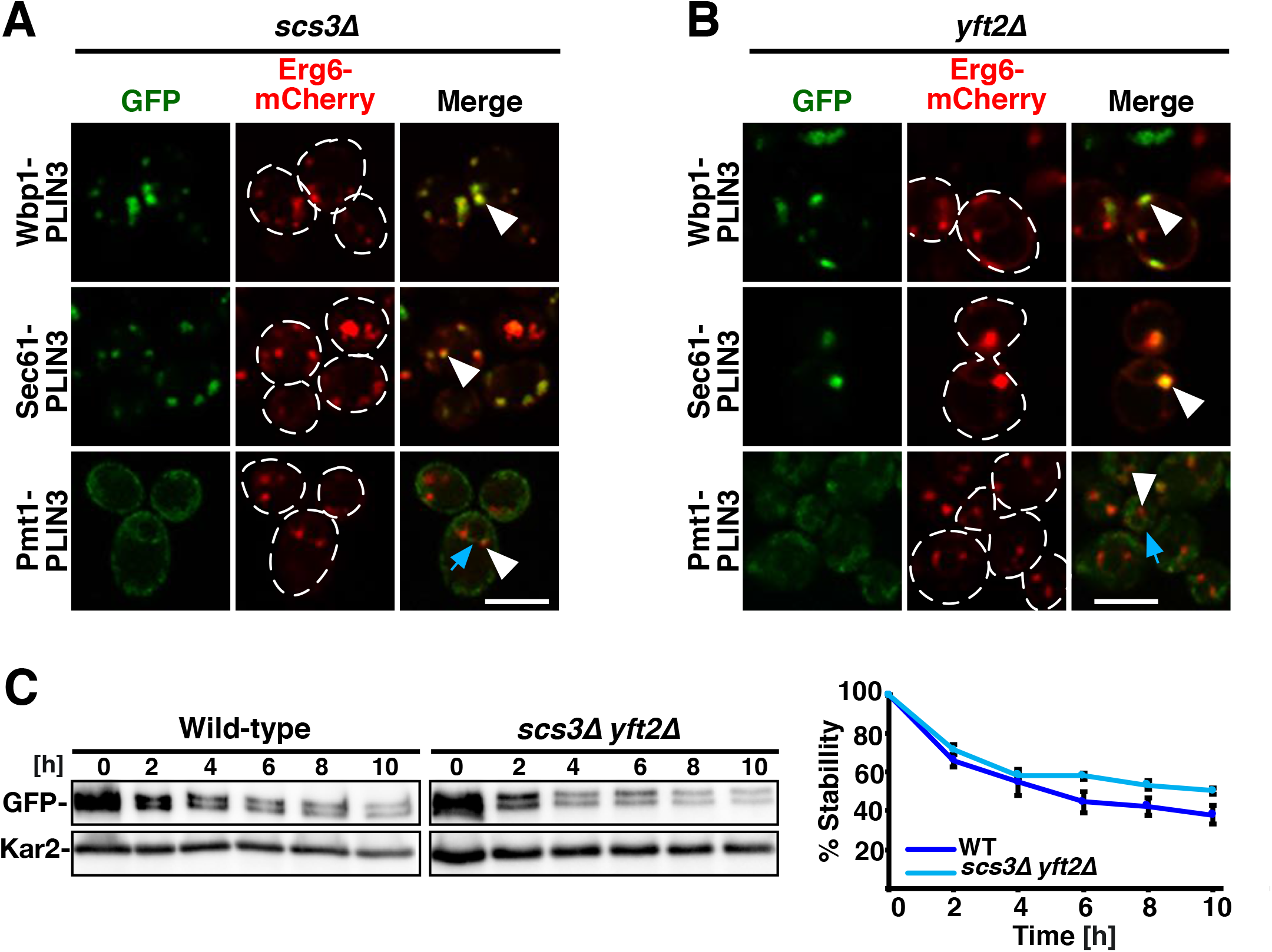
Scs3 and Yft2 share a redundant function in preventing access of Pmt1-GFP-PLIN3 to LDs. A, B) One of the FIT proteins is sufficient to prevent localization of the ER luminal reporter to LDs. Cells lacking one of the FIT proteins, either Scs3 (A) or Yft2 (B) but expressing the indicated membrane-anchored LD reporters together with Erg6-mCherry were cultivated in media containing oleic acid and the localization of the reporters was analyzed by confocal microscopy. White arrowheads indicate punctuate LD localization, blue arrows indicate localization in the ER. Scale bar, 5 μm. C) Stability of Pmt1-GFP-PLIN3 is not affected in wild-type compared to *scs3*Δ *yft2*Δ double mutant cells. Cells were cultivated in oleic acid supplemented medium and poisoned by the addition of cycloheximide (10 μg/ml). Aliquots were removed at the indicated time points and analyzed by Western blotting. The chimeric protein was detected with an antibody against GFP, and detection of Kar2 serves as a loading control. Relative protein stability over time is plotted in the graph.

**Figure S2.**
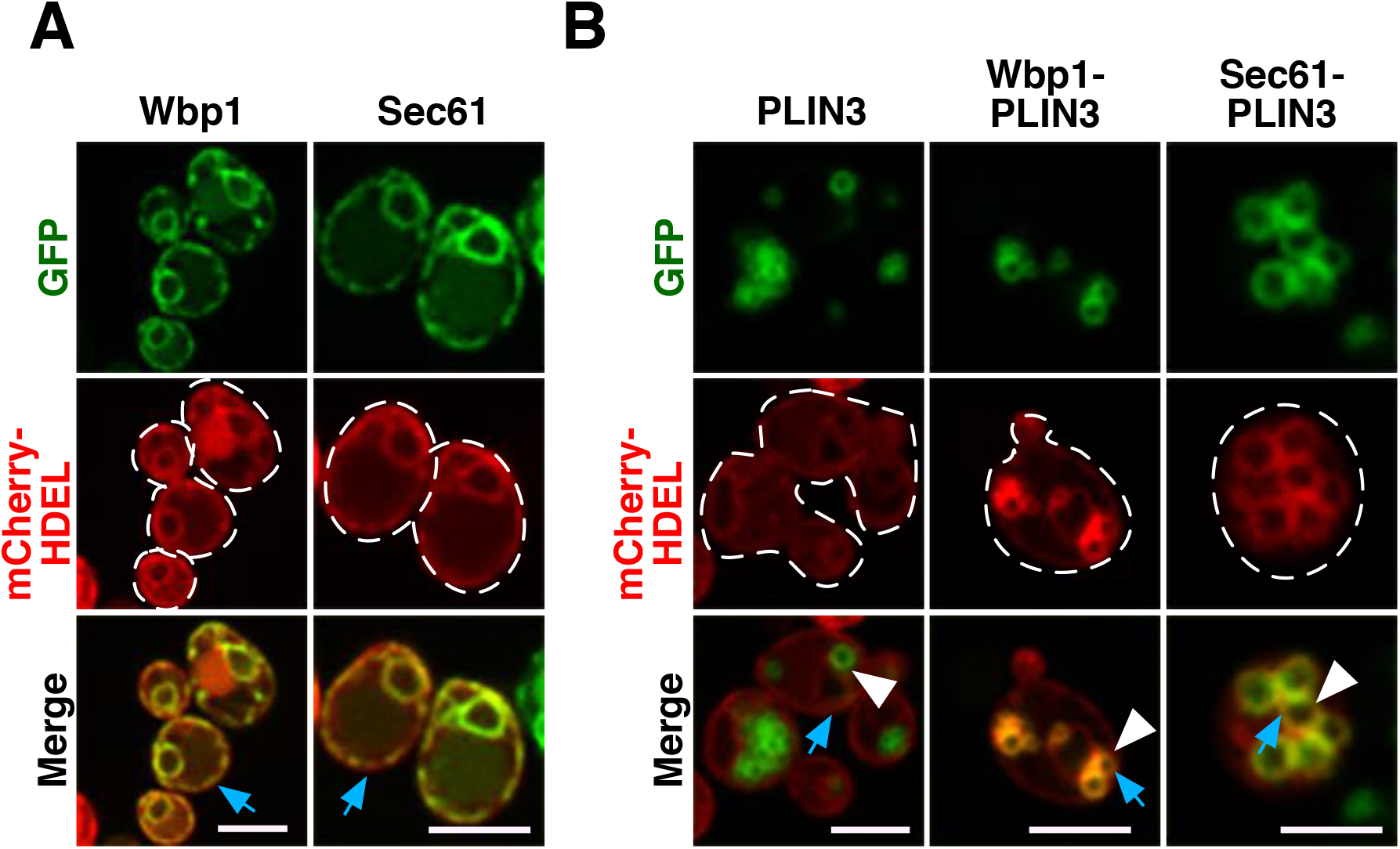
Wbp1-GFP and Sec61-GFP are homogenously distributed within the ER membrane and the LD peripheral localization of Wbp1-GFP-PLIN3 and Sec61-GFP-PLIN colocalizes with the ER. A) Both Wbp1-GFP and Sec61-GFP uniformly stain the ER membrane, colocalize with the ER luminal marker mCherry-HDEL, and do not accumulate at crescent-shaped ER-LD contacts. B) Soluble PLIN3-GFP does not colocalize with the ER marker mCherry-HDEL, but membrane-anchored PLIN3 (Wbp1-GFP-PLIN3 and Sec61-GFP-PLIN3) does. Scale bar, 5 μm.

### Supplementary Tables

**Table S1.**
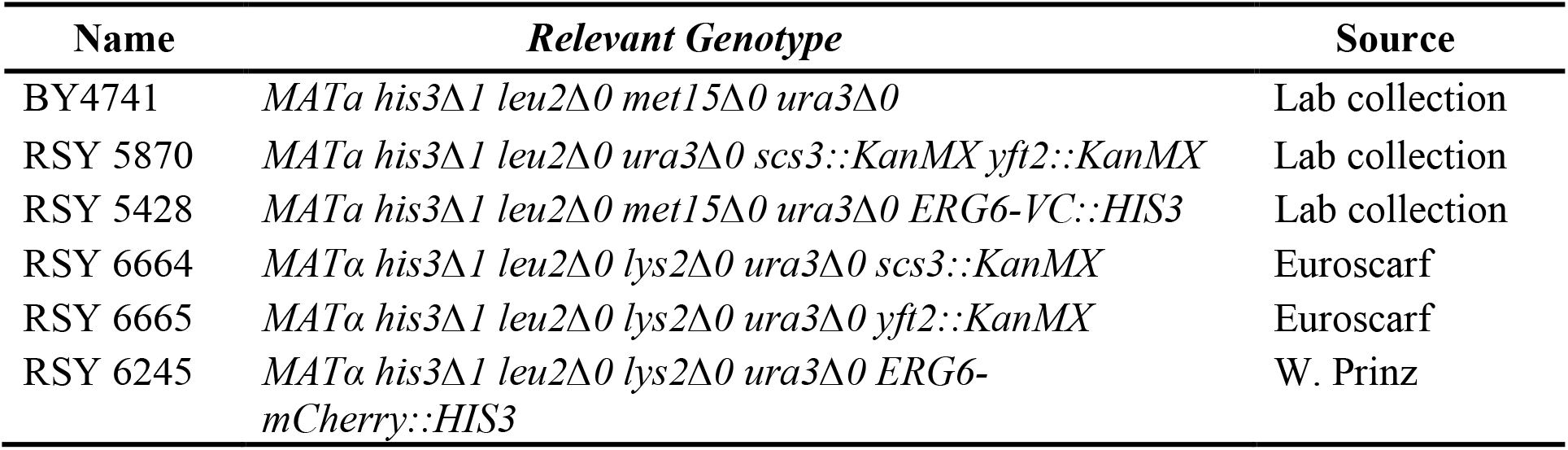
*Saccharomyces cerevisiae* strains used in this study

**Table S2.** Plasmids used in this study

| Plasmids | Source |
| --- | --- |
| pCM189-tetO<sup>7</sup>-PLIN3-GFP/URA3 | This Study |
| pCM189-tetO<sup>7</sup>-WBP1-GFP-PLIN3/URA3 | This Study |
| pCM189-tetO<sup>7</sup>-SEC61-GFP-PLIN3/URA3 | This Study |
| pCM189-tetO<sup>7</sup>-WBP1-mScarlet-PLIN3/URA3 | This Study |
| pCM189-tetO<sup>7</sup>-SEC61-mScarlet-PLIN3/URA3 | This Study |
| pCM189-tetO<sup>7</sup>-PMT1-GFP-PLIN3/URA3 | This Study |
| pCM189-tetO<sup>7</sup>-PMT1-mScarlet-PLIN3/URA3 | This Study |
| pELF1-OST48-GFP-PLIN3 | This Study |
| pCMV-SEC61A1-GFP-PLIN3 | This Study |
| pCMV-POMT1-GFP-PLIN3 | This Study |
| pCM189-tetO<sup>7</sup>-Wbp1-VN/LEU2 | This Study |
| pCM189-tetO<sup>7</sup>-Sec61-VN/LEU2 | This Study |
| pCM189-tetO<sup>7</sup>-ERG6-VC/URA3 | This Study |
| pCM189-tetO<sup>7</sup>-OM14-GFP/URA3 | This Study |
| pCM189-tetO<sup>7</sup>-OM14-GFP-PLIN3/URA3 | This Study |
| pMito-RFP/LEU2 | Nunnari lab |
| pGREG505-ADH-ERG6-mCherry/LEU2 | Lab Collection |
| pRS415-ADH-mCherry-HDEL/LEU2 | Lab Collection |
| pRS416-ERG6-GFP/URA3 | Lab Collection |
| pCMV-mCherry-PLIN2 | Lab Collection |

